# The physical properties of the stick insect pad secretion are independent of body size

**DOI:** 10.1101/2022.03.14.484170

**Authors:** Domna-Maria Kaimaki, Charlotte N. S. Andrew, Andrea E. L. Attipoe, David Labonte

## Abstract

Many insects use adhesive organs to climb. The ability to cling to surfaces is advantageous but is increasingly challenged as animals grow, due to the associated reduction in surface-to-volume ratio. Previous work has demonstrated that some climbing animals overcome this scaling problem by systematically altering the maximum force per area their adhesive pads can sustain; their adhesive organs become more efficient as they grow, an observation which is also of substantial relevance for the design of bio-inspired adhesives. What is the origin of this change in efficiency? In insects, adhesive contact is mediated by a thin film of a liquid, thought to increase adhesive performance via capillary and viscous forces. Here, we use interference reflection microscopy and dewetting experiments to measure the contact angle and dewetting speed of the secretion of pre-tarsal adhesive pads of Indian stick insects, varying in mass by over two orders of magnitude. Neither contact angle nor dewetting speed change significantly with body mass, suggesting that the key physical properties of the pad secretion – its surface tension and viscosity – are size-invariant. Thus, the observed change in pad efficiency is unlikely to arise from systematic changes of the physical properties of the pad secretion; the functional role of the secretion remains unclear.

## 1 Introduction

Bio-inspired adhesives are an important area of biomimetic innovation^1^. However, despite significant progress^2–4^, key challenges remain. Technical adhesives often need to cover large areas and carry heavy loads, resulting in a classic scaling problem: large contacts suffer reduced strength due to stress concentrations, and heavy weights lead to poorer relative performance due to changes in the support-surface-to-volume ratio^4,5^. A similar problem arises for climbing animals of various sizes^6^. How do they resolve this problem?

Previous work has revealed that some climbing animals are able to systematically alter the sustainable force per area – the efficiency of their adhesive pads - as they increase in size^6–10^. In insects^11^, tree frogs^7,8^ and arachnids^12,13^, adhesive surface contacts are mediated via thin films of a liquid secretion, and it has been speculated that systematic changes to the physical properties of this secretion represent a possible strategy to alter pad efficiency with size ^6^. The basis of this hypothesis is the long-standing assumption that the secretion’s functional significance is to promote attachment via capillary and viscous forces^14–19^. Here, we test experimentally whether Indian stick insects alter the physical properties of their pad secretion as they grow, in order to increase pad efficiency and overcome the scaling problem.

## 2 Materials & methods

### 2.1 Study animals

Stick insects (*Carausius morosus*, Sinéty 1901, Phasmatodea, Phasmatidae) were taken from an all-female laboratory colony, fed with bramble and water *ad libitum*, and kept in a climate chamber at 60%RH/25°C with a 12/12 h light cycle. We randomly collected 54 individuals to cover a body mass range of 2.57 – 892.03 mg (EX124, Explorer Analytical Balance, Ohaus, USA; readability = 0.1 mg), close to the maximum mass range of the colony (Fig.1A). All experiments were conducted with live insects and adhesive pads (arolia) on the forelegs, which were confirmed to be free of damage by visual inspection (Leica S APO stereomicroscope, Leica Microsystems, Germany).

**Fig. 1.**
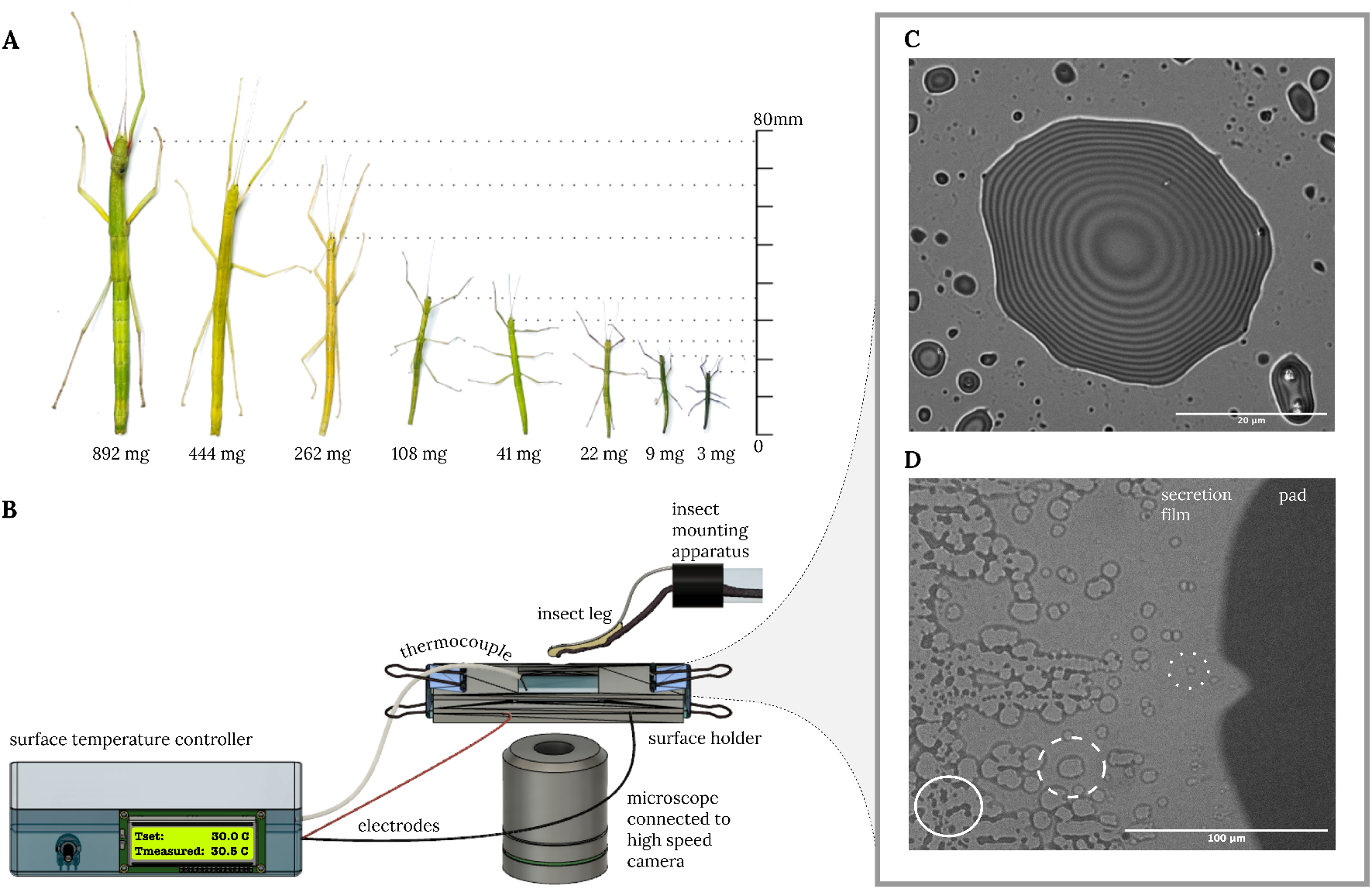
A. *Carausius morosus* stick insects vary by over two orders of magnitude in mass between first instars and adults. B. The physical properties of the secretion were determined with a custom-made setup. C. Interference Reflection Microscopy image of a droplet of the pad secretion deposited on glass, recorded at 30°C. Interference fringes indicate fixed height increments and can thus be used to estimate the contact angle of the pad secretion and hence, its surface tension. D. When a contacting pad is slid to expose the secretion film, thermodynamic instabilities lead to dewetting; dry patches nucleate within the thin continuous film (dotted circle), then grow (dashed circle) and finally merge (full circle). The speed of dry patch growth is inversely proportional to the secretion’s viscosity.

### 2.2 Contact angle and dewetting speed measurements

We estimated the physical properties of the secretion - its surface tension and viscosity - by Interference Reflection Microscopy (IRM) and dewetting experiments, performed with a five-component setup: (i) an insect mounting apparatus as described in Labonte *et al*^20^; (ii) a borosilicate glass coverslip coated with conducting Indium-Tin-Oxide on the bottom side to enable surface temperature control via Joule heating (see SI for surface preparation and characterisation); (iii) a surface holder to secure the coverslip in place and to connect it to a temperature control unit; (iv) a K-thermocouple (TP870, Extech, USA) connected to a custom-built Arduino temperature controller to maintain a constant surface temperature of 30°C during all experiments; and (v) an inverted microscope (DMi8, Leica Microsystems, Germany) connected to a high-speed camera (Blackfly S USB3, BFS-U3-16S2C-CS, FLIR, USA; Fig.1B).

In order to assess whether the secretion’s surface tension changes with size, we measured contact angles of droplets deposited on glass using IRM, following earlier work^21,22^. In brief, we performed artifical ‘steps’ with the mounted arolium, using a micromanipulator (M3301R, World Precision Instruments, UK), which resulted in the deposition of small droplets (average radius ~ 5 *μ*m, below the capillary length). The droplets were then imaged at 63x magnification, an illuminating numerical aperture of 0.8, and with a wavelength of 445 nm from a pE-300 ultra LED (filter with 10 nm bandwidth, CoolLED, UK; Fig.1C). We approximated the contact angle as the slope calculated from the first two height increments derived from the interference fringes, to minimise a bias introduced by the curvature of the droplets. We measured contact angles from three droplets per individual and 23 individuals across the size range (n=23, N=69). The relationship between contact angle and surface tension is complex^23^, but simplifies for a dominantly dispersive fluid such as the insect pad secretion (see SI)^24–26^.

In order to estimate the viscosity of the secretion, we quantified its dewetting speed^22^. Hydrodynamic theory predicts that surface tension, *γ*, and dewetting speed, *v*_d_, are related to viscosity, *η*, via:

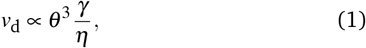

where *θ* is the contact angle^27–29^ (Fig.1D). To quantify dewetting speed, we connected the micromanipulator to a hydraulic drum controller (MHW-3, Narishige, Japan), to rapidly slide pads in surface contact in parallel to the coverslip. The subsequent dewetting of the thin film left behind was recorded at 20x magnification and 226 fps (Fig. 1D). We recorded a minimum of three dewetting videos per stick insect. Every dewetting experiment was conducted on a clean part of the glass coverslip. A period of at least 10 min separated consecutive experiments with the same individual in order to provide time for recovery of a sufficient footprint volume^17^. The rate of growth of dry patches, the dewetting speed (*v*_d_ = *dR/dt*), was extracted with a custom-written ImageJ macro^30^. A minimum of three dry patches, tracked for at least four frames, were analysed for 54 stick insects (n=54, N=354).

## 3 Results & discussion

The contact angle of the pad secretion does not vary significantly with size (log-log slope obtained via ordinary least-squares regression=-0.057 (95% CI (−0.181 | 0.067)), *F*_1,21_ =0.911, p=0.351, n=23; Fig.2A). The mean contact angle of 17 ± 6° (mean ± std), is in excellent agreement with previous measurements in ants, stick insects and cockroaches (18 ±7°, 18 ± 1° and 17 ± 1°, respectively)^17,22,31^. As the secretion is immiscible in water and almost entirely dispersive^24–26^, this result strongly suggests a size-invariant surface tension (see SI for detailed argument).

**Fig. 2.**
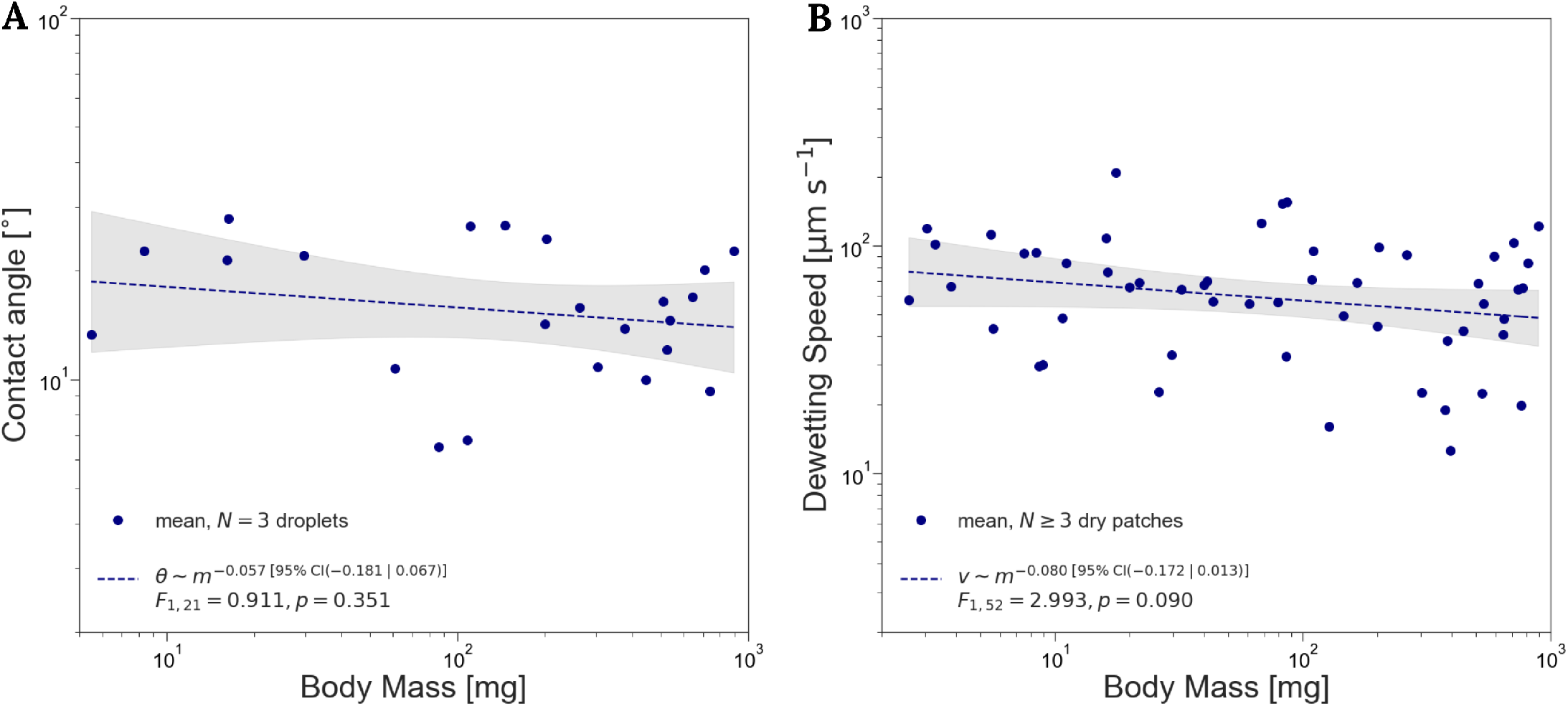
Neither the contact angle (A) nor the dewetting speed (B) of the pad secretion change significantly with size, indicating that both surface tension and viscosity are size-invariant (*n_θ_* = 23 individuals, *n*_dewetting_ = 54 individuals).

The dewetting speed showed a small but insignificant trend to decrease with size (log-log slope=-0.080 (95% CI (−0.172 | 0.013)), *F*_1,52_ = 2.993, p=0.090, n=54; Fig.2B). The mean dewetting speed was 69 ± 39 μms^−1^ (mean ± std), comparable to preliminary results reported for ants (*v*_d_ = 60 ± 5 μms^−1^)^22^. As both contact angle and dewetting speed do not vary significantly with size, neither does viscosity (log-log slope=0.004 (95% CI (−0.149 | 0.158), *F*_1,52_=0.003, p=0.955; see SI). Any change in pad efficiency is thus unlikely to arise from changes of the physical properties of the pad secretion.

Although contact angle and dewetting speed are size-invariant, they scatter notably around the mean (38% and 56% coefficient of variation, respectively). A linear mixed model with random intercept for the 23 paired individuals suggested that the dewetting speed variation arises largely from inter-individual variation (moderate Intraclass Correlation Coefficient (ICC) of 0.56) and thus, that the dewetting measurements are robust. When the variation in contact angle is included, (*v_d_* /*θ*^3^ ∝ *γ/η* as shown in eq.1), the ICC increases to 0.88, highlighting that the increased variation from the non-linear contribution of the contact angle further accentuates the differences amongst individuals (see SI).

The insect pad secretion is a multi-phasic emulsion, consisting of aqueous droplets dispersed in an oily continuous phase^22,25^. IRM imaging of the pad contact area across the size range revealed a qualitatively similar appearance and a similar amount and size of aqueous phase droplets (Fig.S3). In combination with the size-invariance of the contact angle and dewetting speed, this finding strongly suggests that the chemical composition of the pad secretion may also vary little with size.

## 4 Conclusion

Our results demonstrate that stick insects do not systematically alter the physical properties of their pad secretion to avoid a size-related reduction in safety factors. Indeed, a size-specific change in efficiency is only observed in the presence of shear forces^10^, and is thus likely explained by the shear-sensitivity of biological adhesive pads^9,32,33^. Potential changes in the physical properties of the pad secretion will only affect adhesive performance if attachment is dominated by capillary or viscous forces. Although this has been a long-standing assumption^15–19^, strong direct evidence in support of it is absent, and recent research has put forward alternative functional interpretations^34–6^. Systematic changes in adhesive performance may also arise from a size-specific variation of elastocapillary effects^37^, or the pad’s viscoelastic properties, which thus need to be investigated in future work.

## Competing interests

The authors declare that they have no known competing financial interests or personal relationships that could have appeared to influence the work reported in this paper.

## Funding

Funding for this study was provided by the Biotechnology and Biological Sciences Research Council to DL (BBSRC grant no. BMPF-P72408).

## Contributions

D-MK: methodology, investigation, supervision, formal analysis, visualisation and writing - original draft, CNA: investigation, formal analysis and writing - review and editing, AELA: methodology, formal analysis and writing - review and editing, DL: conceptualisation, funding acquisition, supervision, formal analysis and writing - review and editing.

## Acknowledgements

The authors would like to thank Myrta N. Stoukidi and Eleni Papafilippou for their respective contributions to the design of the surface temperature controller and the ImageJ dewetting analysis macro, and Fabian Plum for the 3D scan of the stick insect leg appearing in Fig.1.

## Notes

### Competing Interest Statement

The authors have declared no competing interest.

