## Supplementary Information for "The physical properties of the stick insect pad secretion are independent of body size"

### Surface preparation and characterisation

We used borosilicate glass coverslips coated with conducting Indium-Tin-Oxide (ITO) on one side (18 x 18 mm, #1.5 thickness and $8-12\,\Omega$ resistance; 06481-AB, SPI Supplies, USA). The coverslips were selected to provide the transparency and conductivity necessary to (i) obtain Interference Reflection Microscopy images and (ii) control the surface temperature using Joule heating (supplied by the surface temperature control unit), respectively. The viscosity of liquids is sensitive to temperature. We controlled surface temperature by placing the coverslip on a surface holder, such that the ITO layer was on the bottom, and connected to a K-thermocouple and custom-made Arduino temperature controller (see Fig.1B in main article). We kept the surface temperature at 30°C, high enough to be insensitive to temperature fluctuations in the room, and low enough to be comparable to the environmental chamber which houses our laboratory colony (25°C).

Prior to experiments the coverslips were cleaned in an ultrasonic bath in acetone and isopropanol for 10 mins each, followed by 30 s of rinsing in DI water; surfaces were blow-dried with compressed air. The surface energy of the coverslips was quantified by sessile drop contact angle measurements performed with a custom-made goniometer equipped with a Dino-lite Edge digital microscope (AM7915MZTL, AnMo Electronics Corporation, Taiwan). A pipette was used to dispense 1 µl droplets onto each surface; images of N=2-5 droplets per surface and n=10 surfaces were captured. Contact angles were extracted using the ellipse fit of the contact angle plugin in Fiji 2.1.0/1.53c[1]. Three liquids (diiodomethane, formamide and DI water; Sigma-Aldrich, USA), varying in their proportion of polar and dispersive surface tension, were utilised to quantify the surface energy of the coverslips, $\gamma_S$, by simultaneously solving the Young-Dupré and Owens-Wendt (equations (1) and (2) below); regression was conducted using the polyfit method within NumPy[2] and the statsmodels API[3] in Python v3.7.4[4]. The polar and dispersive components of the surface energy were then determined as the square of the slope and y-intercept of the best fit line, respectively (Table S1).

Table S1 Surface energy results for glass coverslips determined by linear regression of data from three solvents: diiodomethane ($\gamma^p = 0$ mNm$^{-1}$, $\gamma^d = 50.8$ mNm$^{-1}$, $\theta = 41 \pm 5°$), formamide ($\gamma^p = 19.0$ mNm$^{-1}$, $\gamma^d = 39.0$ mNm$^{-1}$, $\theta = 40 \pm 7°$) and DI water ($\gamma^p = 51.0$ mNm$^{-1}$, $\gamma^d = 21.8$ mNm$^{-1}$, $\theta = 52 \pm 9°$).

| | $\theta_{water}$ [°] | $\gamma_S^p$ [mNm$^{-1}$](95%CI) | $\gamma_S^d$ [mNm$^{-1}$](95%CI) | $\gamma_S$ [mNm$^{-1}$](95%CI) | $R^2$ |
|---|---|---|---|---|---|
| Sonicated borosilicate glass | $52 \pm 9$ | 17.3 (10.6\|24.0) | 35.1 (25.8\|44.4) | 52.4 (41.0\|63.9) | 0.97s |

### The relationship between contact angle and surface tension

The equilibrium contact angle a droplet makes when placed on a solid surface is the result of a balance between the forces arising from adhesive and cohesive interactions between and within the three contacting phases, as described by the Young-Dupré equation[5,6] (provided its diameter is below the capillary length):

$$\gamma_S = \gamma_{SL} + \gamma cos\theta \tag{1}$$

Here, $\gamma_S$, $\gamma_{SL}$, $\gamma$ and $\theta$ are the surface free energy of the solid, the solid-liquid interfacial tension, the surface tension of the liquid and the equilibrium contact angle, respectively.

The solid-liquid interfacial tension may be estimated as the difference between the bulk surface free energies and the sum of the geometric means of all dispersive interactions, caused by temporary fluctuations in the charge distribution between molecules, and polar interactions, resulting from hydrogen or covalent bonds or dipole-dipole interactions[7,8]:

$$\gamma_{SL} = \gamma_S + \gamma - 2\left(\sqrt{\gamma_S^d \gamma^d} + \sqrt{\gamma_S^p \gamma^p}\right), \tag{2}$$

Combining equations (1) and (2), eliminates the interfacial tension, so that the contact angle only depends on the dispersive and polar surface energies of the bulk liquid and solid. Thus, the surface energy of a solid may be estimated by performing contact angle measurements with multiple liquids with known polar and dispersive surface tension[9–11] (see previous section). Conversely the polar and dispersive surface tension of a liquid may be estimated by performing contact angle measurements on multiple surfaces with known energy[12]. The insect pad secretion is immiscible in water, and dominantly dispersive with negligible polar components ($\gamma^p \approx 0$) [12]. The surface tension of the liquid thus becomes a sole function of the dispersive surface energy of the solid, and the static contact angle:

$$\gamma \approx \frac{4\gamma_S^d}{(1+cos\theta)^2} \tag{3}$$

Hence, a size-invariant contact angle, measured on the same surface, implies a size-invariant surface tension; using the mean and standard deviation of contact angle as measured (see main manuscript), and $\gamma_S^d = 35.1\ \mathrm{mNm^{-1}}$, we find a surface tension of $37 \pm 9\ \mathrm{mNm^{-1}}$.

### Viscosity estimation

The viscosity of the pad secretion can be estimated using the dewetting equation ($v_\mathrm{d} \propto \theta^3 \frac{\gamma}{\eta}$; see main article).Considering a surface tension of $37 \pm 9\ \mathrm{mNm^{-1}}$ (see previous section), a dewetting speed as shown in Fig.2B (see main article) and a contact angle that was either measured (Fig.2A in the main article; 23 paired individuals) or the mean of the measured (17.3°; 21 unpaired individuals), we calculated the viscosity and showed that it is size-invariant (log-log slope=0.004 (95% CI (-0.149 | 0.158), $F_{1,52}$=0.003, p=0.955; Fig.SS1).

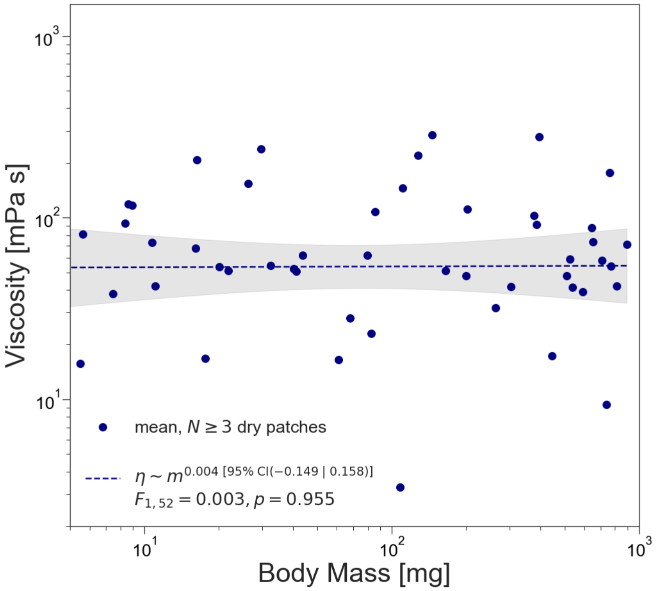


Fig. S1 The viscosity of the pad secretion does not change significantly with size (n=54 individuals).

### Discussion on Intraclass Correlation Coefficient

The coefficient of variation ($CV = \frac{\sigma}{\mu}$) was calculated using data from 23 individuals for the contact angle and 54 individuals for the dewetting speed (data provided in a Supplement). For further analysis, a linear mixed model with a random intercept was implemented using the statsmodels API[3] in Python v3.7.4. The linear mixed model was implemented on the paired data, i.e. data from individuals for which both contact angle and dewetting measurements were recorded (n=23 individuals, N=137 dewetting measurements). The Intraclass Correlation Coefficient[17] was calculated as:

$$ICC = \frac{\sigma^2_{\mathrm{group}}}{\sigma^2_{\mathrm{group}} + \sigma^2_{\mathrm{unexplained}}}, \tag{4}$$

where $\sigma^2_{\mathrm{group}}$ is the variance of the grouped parameter (individual stick insect) and $\sigma^2_{\mathrm{unexplained}}$ is the unexplained variance (denoted as the scale in the statsmodel output). This formula was first applied to the dewetting speed data to assess the inter-individual variation. The resulting ICC is moderate (0.56) suggesting that the variation largely stems from the biological variation across individuals (Fig.S2). In order to take the contact angle variation into account, we performed the same analysis for $v_\mathrm{d}/\theta^3$ (see eq.4). The non-linear contribution of the contact angle resulted in an increased variation and accentuated the differences amongst individuals (strong ICC of 0.88).

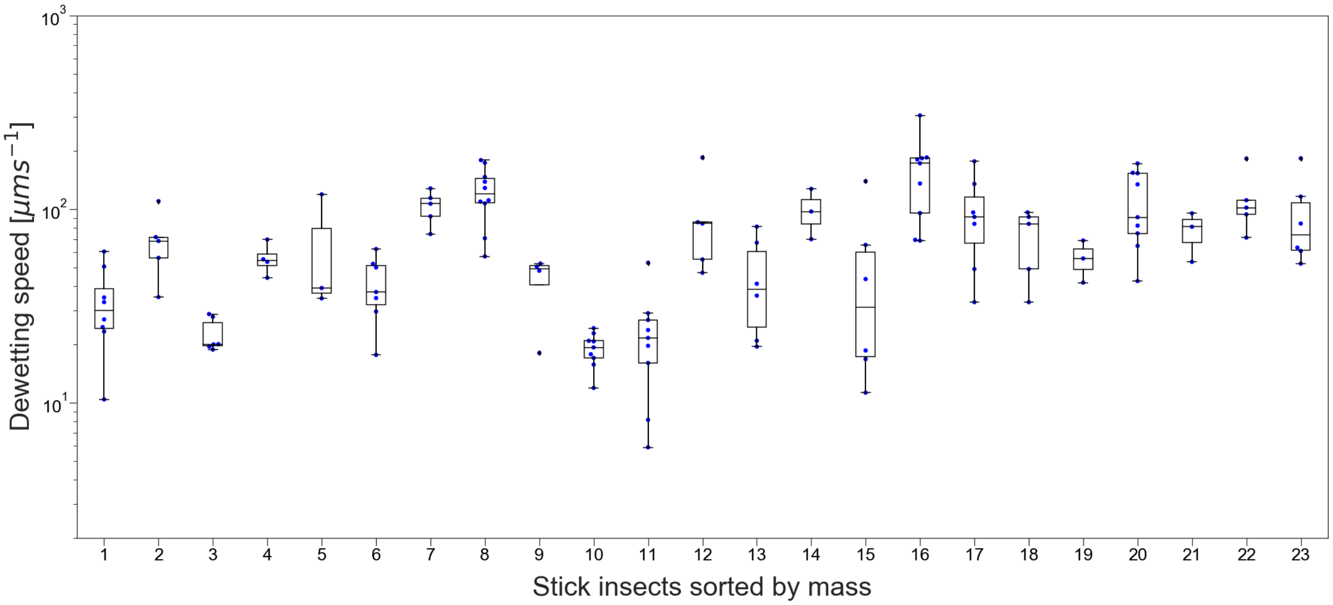


Fig. S2 Variation in dewetting speed and thus viscosity arises largely from inter-individual variation. Only an insignificant amount of variation is explained by body mass.

**Two-phasic composition of the pad secretion**

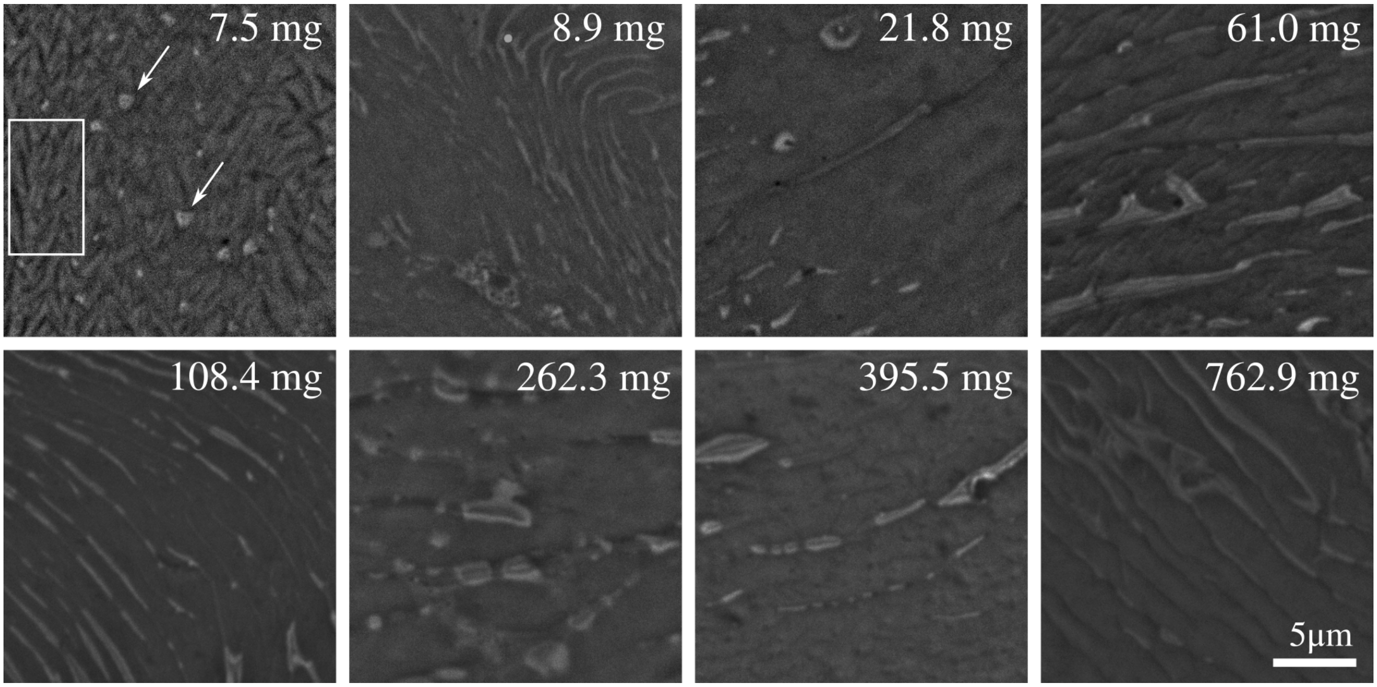


Fig. S3 Interference Reflection Microscopy images of the contact area of adhesive pads of *Carausius morosus* across size following Federle *et al*[18, 19–21]. Areas of high contrast represent the disperse aqueous phase (white arrows). In all instars, the aqueous phase appears in small quantities, and in the form of distinct droplets, often accumulating in cuticular folds (low contrast areas within white box). The qualitative appearance of the contact area is similar across insects of varying size. The absolute amount of the aqueous phase may be larger as the relative area fraction of bright droplets may indicate, as droplets may be of a size below the resolution limit of the light microscope[20]; their small size also renders them extremely volatile, precluding quantitative chemical analysis.

4 van Rossum G. Python Reference Manual; 1995.

5 Young T. III. An essay on the cohesion of fluids. Phil Trans R Soc. 1805;95:65-87.

6 Schrader ME. Young-Dupré Revisited. Langmuir. 1995;11(9):3585-9.

7 Fowkes FM. Attractive Forces At Interfaces. Ind Eng Chem. 1964;56(12):40-52.

8 Owens DK, Wendt RC. Estimation of the Surface Free Energy of Polymers. J Appl Polym Sci. 1969;13:1741-7.

9 Kabza K, Gestwicki JE, McGrath JL. Contact Angle Goniometry as a Tool for Surface Tension Measurements of Solids, Using Zisman Plot Method: A Physical Chemistry Experiment. J Chem Educ. 2000;77(1):63-5.

10 Janssen D, De Palma R, Verlaak S, Heremans P, Dehaen W. Static solvent contact angle measurements, surface free energy and wettability determination of various self-assembled monolayers on silicon dioxide. Thin Solid Films. 2006;515(4):1433-8.

11 Bonn D, Eggers J, Indekeu J, Meunier J, Rolley E. Wetting and spreading. Rev Mod Phys. 2009;81(2).

12 Hasenfuss I. The adhesive devices in larvae of Lepidoptera (Insecta, Pterygota). Zoomorphology. 1999;119(3):143-62.

13 Redon C, Brochard-Wyart F, Rondelez F. Dynamics of Dewetting. Phys Rev Lett. 1991;66(6):715-8.

14 Tanner LH. The spreading of silicone oil drops on horizontal surfaces. J Phys D Appl Phys. 1979;12:1473.

15 Martin P, Brochard-Wyart F. Dewetting At Soft Interfaces. Phys Rev Lett. 1998;80(15):3296-9.

16 Shull KR, Karis TE. Dewetting Dynamics for Large Equilibrium Contact Angels. Langmuir. 1994;10(1):334-9.

17 Koo TK, Li MY. A Guideline of Selecting and Reporting Intraclass Correlation Coefficients for Reliability Research. J Chiropr Med. 2016;15(2):155-63. Available from: `http://dx.doi.org/10.1016/j.jcm.2016.02.012`.

18 Federle W, Riehle M, Curtis ASG, Full RJ. An Integrative Study of Insect Adhesion: Mechanics and Wet Adhesion of Pretarsal Pads in Ants. Integr Comp Biol. 2002;42(6):1100-6.

19 E W MacBride and A Jackson. The Inheritance of Colour in the Stick-Insect, *Carausius Morosus*. R Soc. 1915;89(611):109-18.

20 Dirks JH. Mechanisms of fluid-based adhesion in insects. University of Cambridge; 2009.

21 Peattie AM, Dirks JH, Henriques S, Federle W. Arachnids secrete a fluid over their adhesive pads. PLoS One. 2011;6(5):2-7.
